# Quantitative Label-Free Single-Cell Proteomics on the Orbitrap Astral MS

**DOI:** 10.1101/2024.07.31.605978

**Authors:** Valdemaras Petrosius, Pedro Aragon-Fernandez, Tabiwang N. Arrey, Jakob Woessman, Nil Üresin, Bauke de Boer, Jinyu Su, Benjamin Furtwängler, Hamish Stewart, Eduard Denisov, Johannes Petzoldt, Amelia C. Peterson, Christian Hock, Eugen Damoc, Alexander Makarov, Vlad Zabrouskov, Bo T. Porse, Erwin M. Schoof

## Abstract

Single-cell proteomics by mass spectrometry (scp-MS) holds the potential to provide unprecedented insights into molecular features directly linked to the cellular phenotype, while deconvoluting complex organisms into their basic building blocks. Tailored sample preparation that maximizes the extracted amount of material that is introduced into the mass spectrometer has rapidly propelled the field forward. However, the measured signal is still at the lower edge of detection approaching the sensitivity boundary of current instrumentation. Here, we investigate the capacity of the enhanced sensitivity of the Orbitrap Astral mass spectrometer to facilitate deeper proteome profiles from low-input to single-cell samples. We carry out a comprehensive data acquisition method survey to pinpoint which parameters provide most sensitivity. Furthermore, we explore the quantitative accuracy of the obtained measurements to ensure that the obtained abundances are in line with expected ground truth values. We culminate our technical exploration by generating small datasets from two cultured cell lines and a primary bone marrow sample, to showcase obtainable proteome coverage differences from different source materials. Finally, as a proof of concept we explore protein covariation to showcase how information on known protein complexes is captured inherently in our scp-MS data.

## Introduction

The multicellular intricacies governing human physiology are the result of a landscape of coordinated interactions between trillions of single cells that constitute the human body. With the arrival of single-cell RNA sequencing (scRNA-seq), our understanding about the phenotypic diversity present in cell populations which were once thought to be uniform has rapidly increased^1–3^. However, gene expression alone can not capture the complete molecular context required to characterize the phenotypic state of the cell^4–7^. Thus, modalities expanding across other dimensions of molecules, such as proteins, are necessary to gain deep fundamental understanding on how different cellular states are truly defined.

In the past, measuring protein abundance levels at single-cell resolution was predominantly restricted to methods that relied on affinity or chemical reagents to label a protein of interest, to subsequently estimate its abundance using techniques such as immunofluorescence microscopy, fluorescence activated cell sorting (FACS), or next-generation sequencing (NGS) in conjunction with oligonucleotide-linked reagents^8,9^. In spite of the wide availability of such methods, the direct and systemic quantification of proteins, both at a global scale and at the single-cell level remains challenging.

Over the last three decades, mass spectrometry (MS) has established itself as a powerful analytical tool for the comprehensive characterization of proteins contained in a biological sample. However, only since recent advancements enhancing all aspects of the analytical framework of MS-based proteomics, ranging from sample preparation to data processing^10–15^, have allowed the field to start delving into single-cell applications. Although recent single-cell proteomics by mass spectrometry (scp-MS) approaches are capable of quantifying 1000-2000 protein groups at a moderately applicable throughput^11,16–19^, it is still subject to significant limitations that need to be addressed. Among these, key improvements lie in the number of cells that can be analyzed per unit time and the proteomic depth that can be accurately reached. Studies employing scp-MS have mostly been conducted using Orbitrap (OT) and Time of Flight (TOF) analyzers^16,18,20,21^, however, the implementation of Linear Ion Traps (LIT) for this purpose has recently been successfully demonstrated by us and others^22–24^. Integrating alternative mass analyzers such as LIT highlighted specific advantages that highly sensitive mass analyzers have for scp-MS, albeit that the advantages of LIT are inherently limited by its relatively low resolution^23,25^. A principal obstacle is posed by the need to quantify the miniscule amount of ions that can be produced from a single cell, thereby putting sensitivity at the very forefront of priorities in terms of mass analyzer characteristics. While resolution of OT-based data acquisition remains unsurpassed, extensive ion injection times (IIT) are required, ranging in the hundreds of milliseconds for identification and accurate quantification of single-cell signal^18,26–28^, underlining the need for mass analyzers that strike an optimal balance between sensitivity and high-resolution resolving power.

Here, we aim to gauge the capacity of the Asymmetric track lossless analyser (ASTRAL)^29^ to provide quantitative proteomic profiles from low-input, all the way down to single-cell samples. We first carry out a comparison to Orbitrap-only method performance and follow up by evaluating if previously reported approaches to maximize instrument sensitivity, such as wide window acquisition and use of FAIMS are applicable^30–32^. We then carry out a comprehensive data acquisition parameter optimization to pinpoint the boundaries for optimal parameters for low input and single-cell sample analysis, and benchmark their quantitative accuracy. We conclude our study by profiling the single-cell proteomes of three distinct cell types and showcase how covariation of known protein complexes is quantified.

## 1. Results

### Augmented sensitivity of the Astral analyzer facilitates the identification of low-abundant peptides

To measure the enhancement in instrument sensitivity, we first analyzed 1ng of bulk Hela peptide digest with data acquisition parameters that have been determined previously for both Orbitrap Eclipse and Orbitrap Astral instruments^32,33^. While using the same chromatographic setup, the Orbitrap Astral mass spectrometer was able to quantify almost double the amount of peptides relative to the Orbitrap Eclipse (**Figure 1a**). The coefficient of variation (CV) distribution was slightly lower on MS1 (Orbitrap) level, indicating that the increased number of identifications did not have a negative impact on the overall precision of the measurements (**Figure 1b**). Moreover, the ASTRAL analyzer facilitated the capture of peptides in the lower half of the intensity range, rendering those less abundant peptides in the sample accessible to identification, in line with the higher sensitivity of the Astral analyzer (**Figure 1c**).

**Figure 1.**
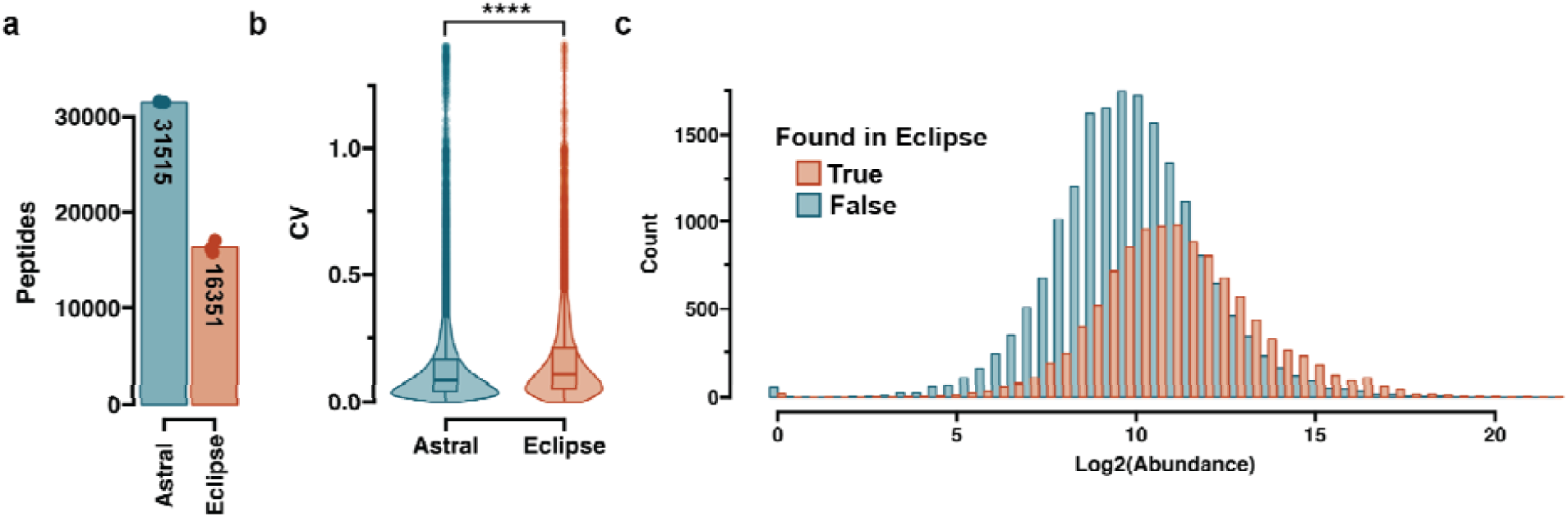
Evaluating the performance of Orbitrap Astral for limited input proteomics. **a)** Identified peptide amount (n=3) at 65 samples per day throughput. Number notes the average number of peptides found **b)** Violin plots of the CVs on non-normalized peptide abundances on MS1 level. T-test was used to assess the significance between the CV distributions. **** notes a p-value ≤ 0.0001. **c)** Histogram showing the identified peptide number relative to the log2 transformed peptide abundances.

#### FAIMS improved proteome coverage of subnanogram samples

The High field asymmetric waveform ion mobility spectrometry (FAIMS) interface has previously been shown to improve the achievable proteome depth by increasing the signal-to-noise of low input samples^30^. However high sensitivity for single-cell samples has recently been demonstrated using narrow window DIA (nDIA)^34^, and without the use of an ion mobility device. This contradicts two previous studies that both utilized ion mobility and reported that wide window isolation can provide significant benefits in sensitivity on Orbitrap-based platforms^31,32^. To investigate this ambiguity, we used a dilution series ranging from 10ng to 250pg and analyzed these samples with a panel of methods where the MS2 scan isolation window is increased to allow for longer injection time while keeping the scan cycle time constant ensuring equal elution peak sampling density (Supplemental Figure 1a-b). Furthermore, we carried out these experiments either with or without the FAIMS interface present on the instrument, to test both aspects simultaneously.

Without the FAIMS device we could see a difference in optimal method for each distinct input level, however no marked overall improvement to relative proteome coverage was observed irrespective of data acquisition settings (Figure 2a). However, contrasting results were obtained when the FAIMS interface was connected. Applying the wide window isolation strategy increased the proteome coverage for all input levels, excluding the high-load samples (Figure 2a). Especially for 250pg the samples, the highest injection time yielded 3-fold higher proteome coverage compared to the lowest, while no such increase was observed without FAIMS. Comparing the overall proteome coverage of high-load samples revealed that the obtainable proteome coverage was comparable with or without FAIMS (Figure 2b), however especially once the input amount reaches below 1ng, FAIMS facilitated a higher identification rate. The technical precision of the common peptides between all methods was similar on the MS2 level for 250pg input, while lower CV values were obtained with FAIMS on MS1 (Figure 2c). Together, these results underline that for high-load samples ≥10ng one can achieve reminiscent performance independent of FAIMS being part of the instrument configuration. However, once the target samples are <10ng, ion mobility can further enhance the sensitivity of low-input proteomics on the Orbitrap Astral MS.

**Figure 2.**
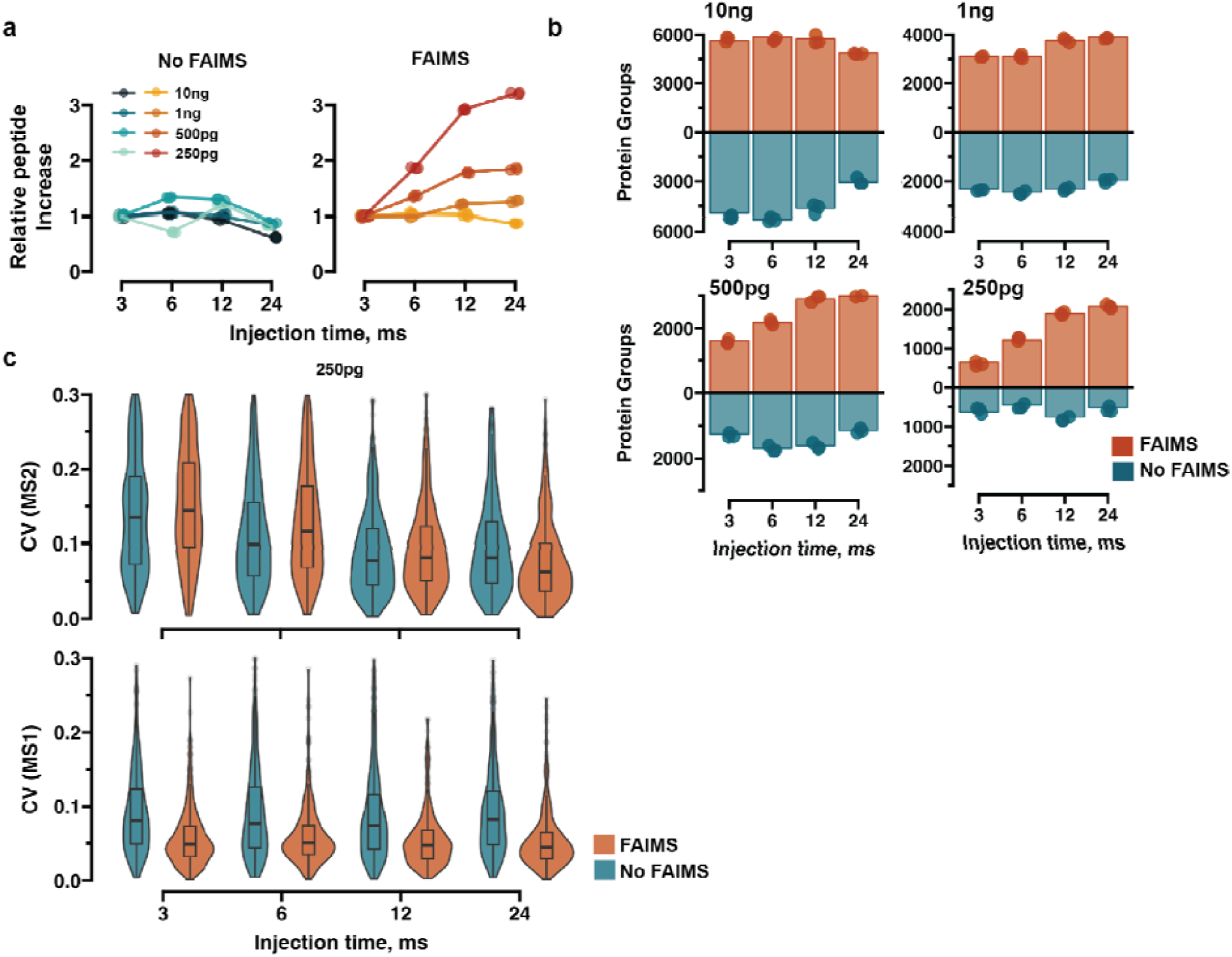
Assessing the utility of FAIMS for Orbitrap Astral with wide-window DIA. **a)** Scatter plot showing the relative changes of proteome coverage with and without FAIMS with different DIA acquisitions methods where injection time on MS2 level is increased and the scan cycle time is compensated by doubling the isolation window. The shortest injection time (3ms) is set as the control value in all cases **b)** Barplot showing the absolute proteome coverage related to **a).** Bar height represents mean proteome coverage and the points individual measurements **c)** Violin plots of CVs on MS1 and MS2 levels for the 250pg sample.

#### Optimizing MS1-based DIA acquisition method for low input samples

Our previously developed high sensitivity, wide isolation window, MS1 quantification based DIA method (WISH-DIA) was tailored to an Orbitrap-only platform^32^, so we aimed to adapt it to efficiently harness the power of the Astral analyzer as well. First we generated a panel of methods where different injection times for the MS2 scans were used, while the MS1 injection time was held constant. We observed that 30-40ms ion injection time yields the best coverage, with 3,978 protein groups and 20,002 peptides being detected from just 250pg input (Figure 3a). We then tested higher MS1 injection times to determine it would provide further coverage improvements, however we did not observe notable increases in achieved proteome depth (Figure 3b). We hypothesized that although we did not gain identifications, we could expect higher quality of quantification due to better ion counting statistics. To test this, we next analyzed a peptide dilution series spanning from 120-1,000pg. With both 100 and 200 ms injection times, the overall measurement error distributions, which was calculated as the percentile difference between the expected and observed value (see Methods), were highly comparable (Supplemental Figure 2a). Furthermore, in the cumulative distribution function (CDF) plots we could see that the overall cumulative distribution of the absolute errors was highly similar, with almost identical fractions of peptides identified 10 and 20 % absolute error (Supplemental Figure 2b).

**Figure 3.**
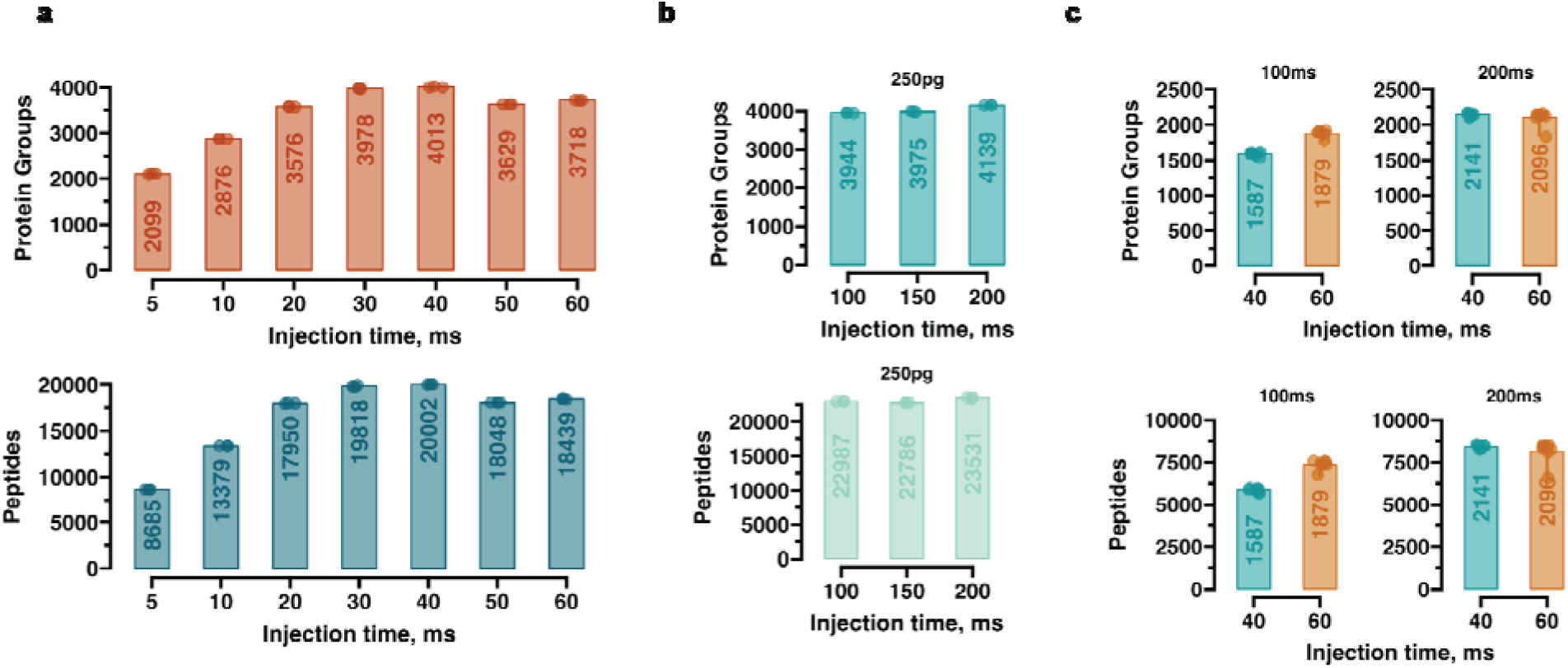
Optimizing Astral parameters for quantitative limited-input proteomics. **a)** Barplot showing the protein groups and peptides identified for 250pg input with different MS2 (Astral) injection times. **b)** Similar to **a)** but with different injection times on the MS1 (Orbitrap) level. **c)** Method optimization for single-cell input. Barplots show identified protein groups and peptides with different injection times on MS2 (40 and 60 ms) and MS1 (100 and 200 ms). All runs carried out at a throughput of 65 samples per day (∼22 min run-to-run).

We then utilized single-cell HEK293 samples that have been prepared with a FACS-based 384-well plate workflow^19^ to determine if the parameters would also yield good performance for actual single-cell samples. We tested the two MS2 injection times of 40 and 60 ms and two MS1 injection times of 100 and 200 ms, to see if the additional ion injection time could uncover more protein groups from such samples. The highest MS2 injection time of 60ms was superior to 40ms, and the best results were obtained with methods that used 200ms injection time at MS1 level (Figure 3c). While for peptide dilutions, 100ms appeared to be the optimal method, for real single-cell samples, using 200ms appears worthwhile as it increases the proteome depth (∼15%) for these samples.

#### Comparison of MS1 and MS2 level quantification for low input proteomics

During MS1 scans, the co-eluting background ions can interfere with the precursor peaks, in turn negatively impacting the accuracy of the measured peak area. Upon isolation and fragmentation for subsequent MS2 scans, these co-isolation effects are deconvoluted, leading to MS2 quantification often being the method of choice in DIA-based proteomics^34^. To gauge the quantitative performance of MS1 and MS2 level peptide ion signal intensities at loads that are reminiscent of ultra-low-input samples, we used 50pg of peptide digest as our reference and compared how accurately we could reconstitute the expected peptide abundance measurements with different target samples spanning a 2 to 10 dynamic range (Figure 4a). Irrespective of quantification method, we could observe a widening of the relative error distribution, indicating decreasing accuracy (Figure 4a-c) which one would expect with the concurrent increase in dynamic range of the ratio. The peak width remained narrower with fragment-based measurements (MS2), albeit MS1 level quantification seems to provide a higher number of peptide identifications with error below 25% for the smallest evaluated ratios (Figure 4b-c).These results corroborated our previous findings that both MS1 and MS2 quantification can be applied when working with low-input samples.

**Figure 4.**
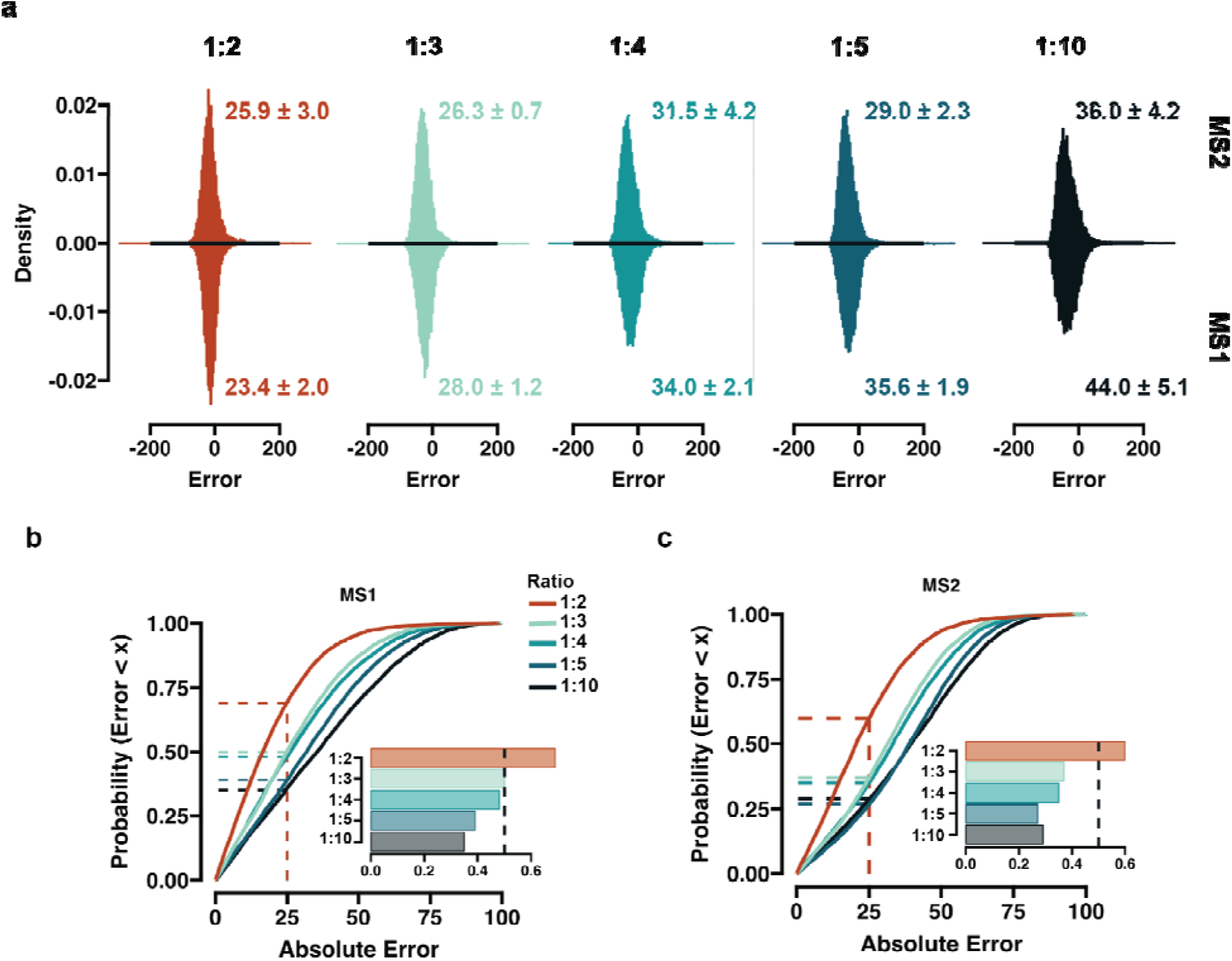
Accuracy comparison of MS1 and MS2 level based quantification. **a)** Histograms showing the relative error distribution on MS2 (top) and MS1 (bottom) level. 50 pg is set as reference, and the ratios correspond to comparison with 100, 150, 200, 250 and 500pg. The numbers note the mean standard deviation of the error distribution from three replicates. **b-c)** Cumulative density function plot showing the absolute error trend for different ratios. Embedded bar plot shows the fraction of peptides below and absolute 25% error. **c)** Histogram showing the distribution of the error on both levels with different ratios. DIA-NN was used for obtaining the peptide abundances.

#### Obtaining quantitative information at approximated single-cell level signal intensities

Currently, it seems common practice in the field that 250pg injections of complex lysates can be utilized as a proxy for actual single-cell samples. In fact, extensive observations in our laboratory over the past years indicate that the amount of actual single-cell derived peptides that reach the instrument is in practice substantially lower^27,35–37^. For this reason, we tested the ability of the Orbitrap Astral MS to provide quantitative measurements at sample loads < 100pg, where we approximate our single-cell sample signal intensity from a 384-well plate based preparation usually lies. We used 100pg as a reference sample and 50, 30, 20 and 10 pg as targets. We excluded any peptides that were not found in all the dilutions and evaluated precision and accuracy for the common list. The median precision was ∼4% in the highest dilutions, which gradually increased, but still remained ∼10% median at 10pg input at peptide level (Figure 5a), showing high reproducibility of measurements of approximated single-cell signal ranges. By examining the overall peptide intensities, we observed downward scaling, however the median value was decreasing more rapidly than expected (Figure 5b). This bias could be due to small losses on the path from the sample vial to the instrument, convoluting accuracy measurements across different dynamic ranges. Despite this, we observed high accuracy with > 70% of the identified peptides carrying a relative error of below 20% (Figure 5c). However, the error distribution widened rapidly with decreasing input amounts and at the lowest dilution, the measurement accuracy became rather questionable (Figure 5d), pointing towards the potential minimal boundary input value for a complex proteome sample. We noted previously that the accuracy and measured dynamic range is negatively correlated (Figure 4). To correct for this, we compared the quantification between our two lowest dilutions and saw the overall error distribution being drastically reduced (Figure 5e). Overall, these findings demonstrate that quantitative measurements are obtainable from single-cell level signals.

**Figure 5.**
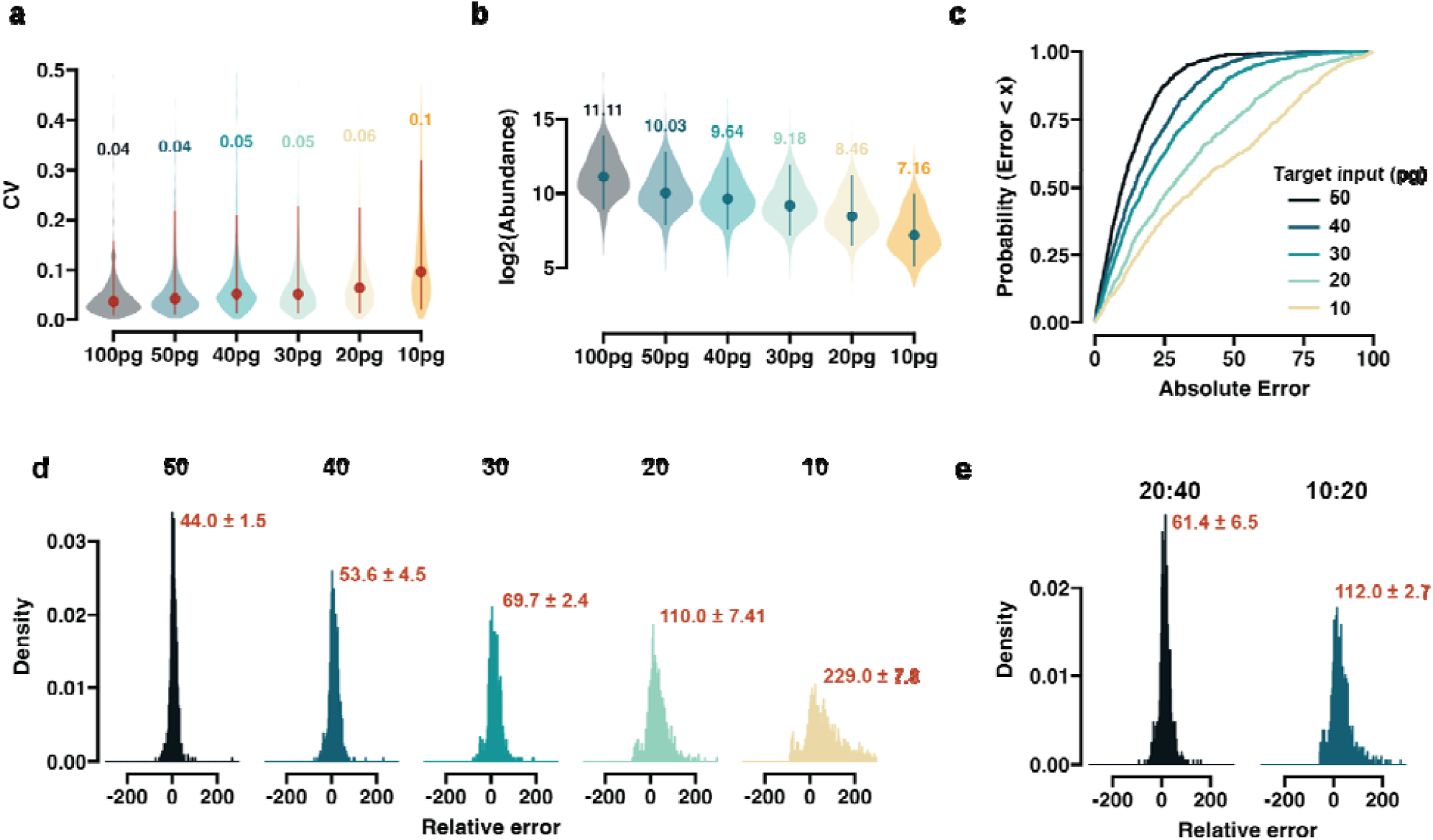
Evaluating the peptide ID propagation quantitative accuracy. **a)**. Violin plot of peptide level CV distributions with different input levels. Only the peptides that are found in all the loads with no missing values are used (n=1064) **b)** Violin plot of peptide log2 transformed abundances. Median value is indicated by the dot and the lines represent the 0.05 and 0.95 quantile boundaries. Numbers note the numeric median value. For both **c)** and **d)** 100 pg is set as reference for the other inputs (50, 40, 30, 20 and 10 pg) as relative targets. **c)** CDF plots of the absolute error. **d) H**istograms showing the relative error distribution based on MS1. The numbers denote the peaked full-width at half maximum (2.634σ) with the uncertainty calculated from three replicates. Replicate 2 data is shown. **e)** Error distribution the same a in **d**, but for 20:40 pg and 10:20 pg comparisons. Spectronaut was used to obtain the peptide abundances.

#### High-Load library effects on quantitative accuracy of the dataset

Higher load libraries consisting of small cell populations or diluted bulk samples have been used to increase the amount of peptide identifications from low-input samples^16,37,38^. Since these peptides are potentially too low in abundance to identify without prior information, we wanted to evaluate the accuracy of these computationally augmented quantifications (Figure 6a). We performed individual searches where 50 pg samples (the ‘reference’ samples) were searched with and without ID propagation between raw files. In individual reference samples, ∼5,000 peptides could be identified (Figure 6b), but if IDs were propagated between all the 50pg runs, we could increase this number by 49%. Including all the files (50 to 500pg) further increases the identified peptides, as including higher input samples into the search reaches three times the number of peptides (∼15000)(Figure 6b), however at the risk of severely affecting quantitative performance.

**Figure 6.**
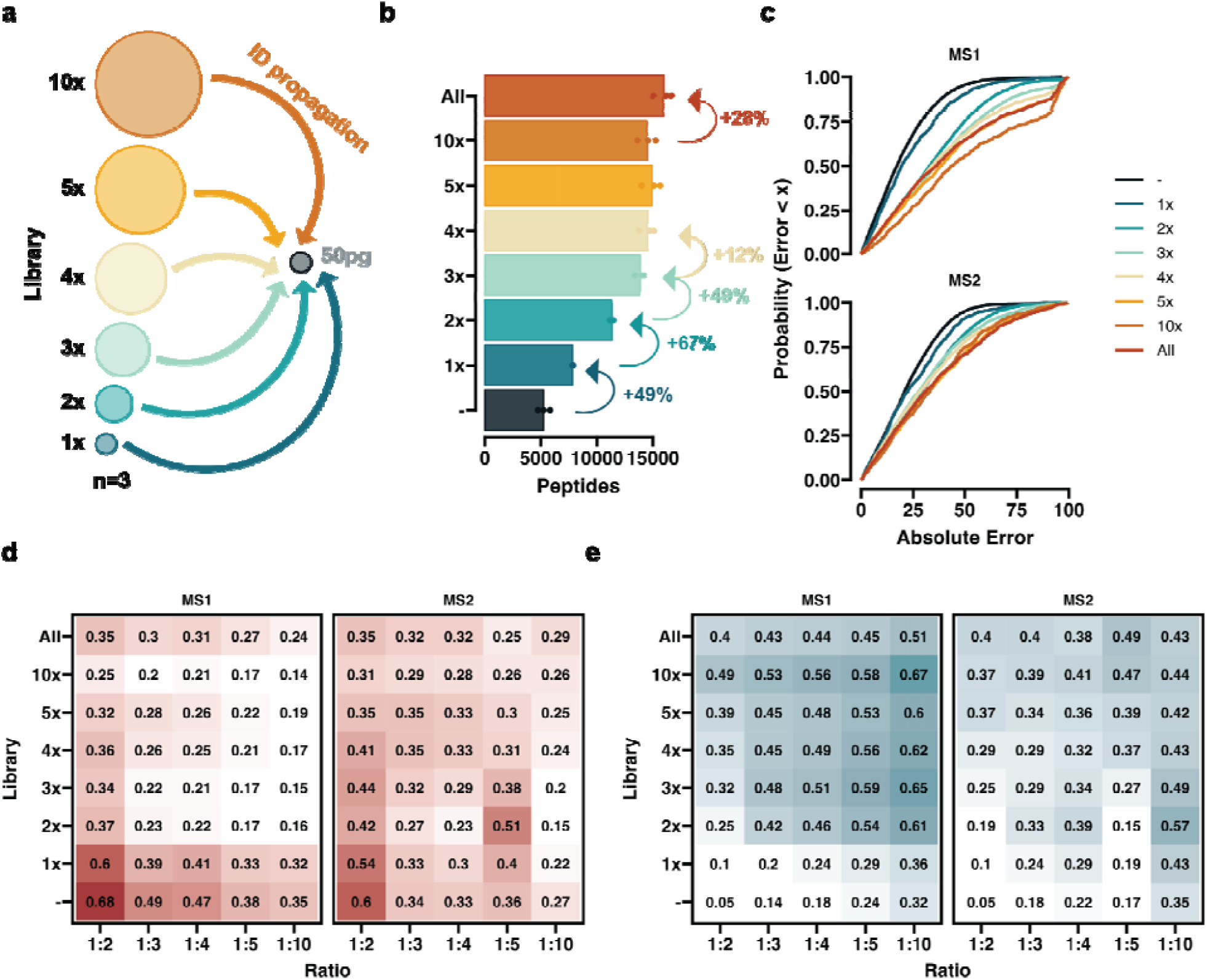
Evaluating the peptide ID propagation quantitative accuracy. **a)**. Schematic depicting the ID propagation to the 50pg reference samples from libraries from different inputs with respect to the reference. **b)** Barplot of the total number of peptides identified without and without libraries. Arrows and numbers denote the fractional increase in peptide identifications. **c)** A CDF plot of the absolute error for the 1:2 comparison (50 : 100 pg). **d-e)** Tile plots showing the fraction of peptide identifications below 25% error **d)** and above 50% **e).** Numbers represent the fraction from the total number of peptides added and the color represents the value size. The same data is used as for Figure 4. Spectronaut was used to obtain the peptide abundances.

To assess the quantitative accuracy of this approach, we used the longest peptide list and annotated the peptides depending on what library size they were first identified in. By plotting the absolute error CDF plots, we could clearly see that peptides found by direct means or horizontal (50 pg to 50pg) ID transfer had lower absolute error compared to peptides propagated from a higher injection amount (Figure 6c). There was a visible drop in MS1 quantification accuracy once peptides gained above 1x library were included, however on MS2 level the difference was not as pronounced. We stratified the peptides into three tiers below 25% (I), between 25-50% (II) and above 50% (III) absolute error. We first examined what peptides added by the libraries were classified as tier I (Figure 6d). With direct or horizontal ID propagation, we see that above 60% of the peptides are in the highest quality category, but this fraction drops as the reference ratio increases, in line with our previous results. Peptides that were identified with the aid of a 2x or higher library not only had a lower fraction of tier I peptides, but more concerningly, an ever growing fraction of lowest quality peptides (Figure 6e). Overall, these results indicate that while the overall number of identifications is enhanced, careful assessment of resulting identification increases are required to extract valuable peptide information and avoid dataset convolution with large numbers of noisy measurements.

#### Quantitative proteome measurements from different single-cell types

The field of scp-MS is still in an extensive state of technical development, with a wide range of experimental assays available on varying instrument platforms. An additional confounding variable is the type of cell used to benchmark the various workflows, making cross-experimental comparisons difficult to achieve. Given the difference in cell size and proteome content / complexity of various cell types, one would expect the achievable proteome coverage to vary substantially depending on the cell population under investigation. To test this, we selected human embryonic kidney cells (HEK293), monocytes (U937) and primary human CD34+ bone marrow (BM) cells, and processed these cells with our standard 384-well plate based workflow^32^. We could identify ∼3,500 protein groups from HEK293 cells, ∼2,500 protein groups from U937 cells and around 1,300 protein groups from primary CD34+ BM cells (Figure 7a). Although both HEK293 and U937 cells are similar in size, we saw a 30% drop in coverage, which could potentially be attributed to the more specific proteome expressed in monocytes relative to HEK293. Expectedly, the identification numbers further decreased in samples generated from primary material, as the very primitive CD34+ BM hematopoietic cells are expected to have more specialized proteomes (to support their multipotency), but also are 3-4 times smaller in size. It has been previously noted that cell size within a cell type is correlated to the number of identified protein groups as more signal can be gained from larger cells^11^. In line with this, we saw a linear correlation between cell size and median sample intensity, which led to higher coverage (Figure 7b).

**Figure 7.**
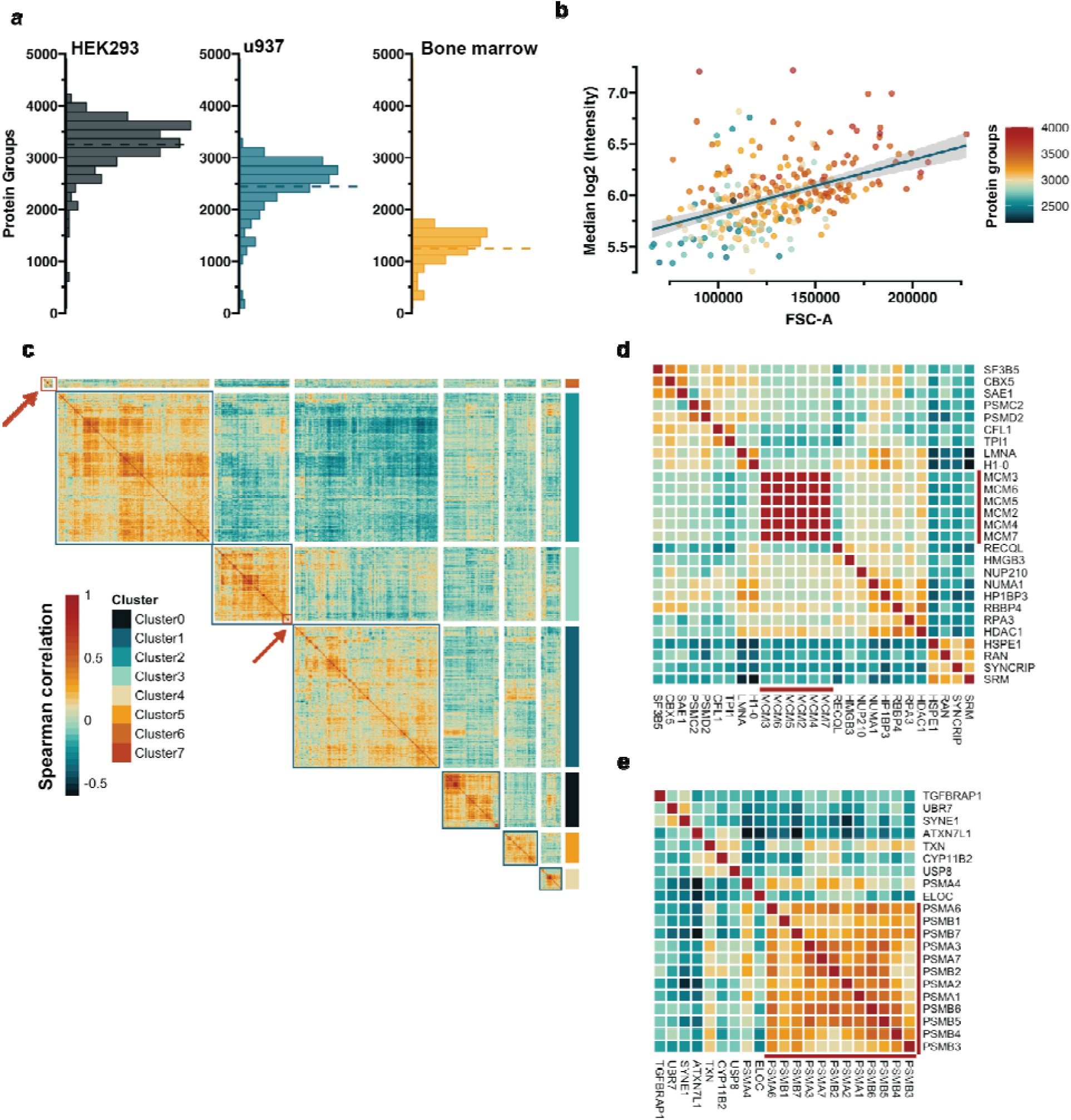
Profiling protein covariation at single-cell resolution. **a)**. Histograms showing the distribution of protein groups obtained from different types of cells HEK293 culture embryonic kidney cells, U937 culture monocytes and primary BM CD34+ cells **b)** Scatter plot of single-cell sample intensity (HEK293) dependency on the isolated cell size. Forward side scatter area (FSC-A) is shown on the x-axis, and the y-axis reflects the median log2 transformed sample intensity. The color denotes the number of protein groups identified in each cell and the line is a linear fit of the data with the confidence interval in grey shade. **c)** Upper diagonal correlation of map of proteins from the HEK293 cells. The map is ordered by hierarchical clustering and stratified into specific groups based on the dendrogram (not shown). Color represents spearman correlation value. Red squares and arrows note the section used to create **d** and **e. d-e)** Cutouts of the correlation map showing the capture covariation of the MCM complex and the proteasome subunits.

To evaluate whether the data captures biologically relevant information, we performed a covariation analysis on the HEK293 dataset. We calculated the protein-wise spearman correlation and filtered out the proteins with no markedly correlated proteins, and subsequently visualized the results with a clustermap (Figure 7c). We could stratify the proteins based on their correlation into modules with hierarchical clustering. We here chose a small number of clusters, but visually evaluating the modules, one can observe the presence of higher granularity, indicating that a higher number could be selected for a more detailed exploration. To showcase specific examples, we selected two small sections from the clustermap, showing the high association of the MCM helicase complex subunits and proteasome subunits (Figure 7d-e), supporting that known biological trends can in fact be quantified.

## Discussion

In this study, we explore the quantitative accuracy of peptide- and protein-level measurements obtained from approximated single-cell level signal intensity and showcase that valid measurements can be obtained from such input. We carry out a series of benchmarking and optimization experiments to adopt previous practices that maximize the sensitivity of the Orbitrap Astral mass spectrometer and resolve some emerging dichotomy between approaches (Figure 2). We proceed to identify optimal parameters for low-input and single-cell samples at a fixed chromatographic gradient and assess the quantitative accuracy of the measurements obtained signal intensities reminiscent of single-cell samples (Figure 2-5). We conclude our study by generating small single-cell datasets and highlight the differences in expected proteome coverage based on cell type and showcase the ability to recapitulate prior biological information (Figure 7).

Single-cell proteomics is advancing at an incredible rate, with especially label-free approaches having garnered a lot of attention and development lately due to the simplicity of such experimental assays^16,31,32,35,37,39^. Label-free methods have achieved the largest reported proteome coverage of single cells by a large margin compared to multiplexing, but at a significant trade-off in throughput. While multiplexed approaches can analyze hundreds of cells per day^19,40,41^, label-free is generally limited to 20-50 samples-per-day (SPD), with some showcases of ∼70-100 cells per day^16,31,32,35,37,42^. DIA-compatible multiplexing approaches could strike a balance between the best features of label-free and multiplexed, but additional efforts are needed to offset spectral complexity, distortion of quantification across channels, and labeling inefficiency issues ^43–45^. Additionally, to further advance label-free approaches, additional improvements in chromatography are required to potentially better harness the power of next-generation instrumentation.

The race towards the greatest depths of the single-cell proteome appears to have taken center stage, with the quantitative accuracy of the obtained measurements receiving far less spotlight. Assessing this is challenging as peptide extraction and inherent heterogeneity of a cell can drastically bias accuracy measurements, underlined by large deviations from the expected values^38,46^. Here, we specifically focused on low pg samples, where the signal intensity is reminiscent of single-cell samples, to assess the quantitative quality of the obtained data, and reflect the capabilities of the instrument at the extreme low end of the sensitivity spectrum (Figure 4-6). In the range of 50-200 pg, the majority of the peptides were measured with an error of below 20% (Figure 4) and a large portion of identified peptides remained below this threshold when the input was decreased to 20pg (Figure 5). Taken together, this indicates that the ASTRAL is capable of providing accurate peptide abundance values from signal intensities that would be expected from single cells.

DIA search augmentation with higher load spectral libraries has been used previously to extend the number of identified protein groups^37,39,47,48^, however given the potentially low quality of the signals, it remains questionable if all the additional values could be considered equally quantitatively accurate. We stratified the peptides based on their input level that allowed their identification and examined the fraction of peptides with a low <25% and high > 50% error (Figure 6d-e). Although we found that with higher-load libraries, a moderate fraction of the peptides were in the lowest error bin, computational approaches to filter library-propagated peptides that deviate too far from the true quantitative values are needed.

Finally, with our optimized method we generated three small datasets of HEK293, U937 and primary CD34+ bone marrow cells (Figure 7). We showed the obtained proteome coverage to be dependent not only on the cell size, but cell type as well (Figure 7a-b). Although the correlation to cell size has been noted previously, this clear distinction between different cell types has not previously been demonstrated. To validate that the generated proteome profiles contain biologically meaningful information, we carried out a covariation analysis on the HEK293 dataset. We visualized the protein-wise correlation values in a clustermap and highlighted two functionally very different complexes where known protein subunits formed tight clusters (Figure 7c-e). When taken together, this work underlines the impact of the increased sensitivity and speed of the ASTRAL mass analyzer on our ability to advance our understanding of cell state heterogeneity at the proteome level.

## 1. Methods

### Experimental Design and Statistical Rationale

The aim of the study was to assess performance of the Orbitrap Astral mass spectrometer for low-input and single-cell samples. To this aim, a variety of different data acquisition methods were used to analyse different amounts of input sample. Experiments that used diluted peptide digest were carried out in triplicate and for single-cell runs at least 8 cells were run per method. The chosen higher number of replicates for single-cells was to account for heterogeneity in the input material. Statistical tests were not used to assess differences in performance between different workflow replicates.

### Cell culture and FACS sorting

HEK cells were cultured in RPMI media containing 10 % FBS and 1 % Penstrep. Upon 80% confluence, cells were harvested and washed three times with ice-cold PBS to remove any remaining growth media prior to cell sorting.

Cell sorting was done on a FACS Aria III or Aria II instrument, controlled by the DIVA software package (v.8.0.2) and operated with a 100 μm nozzle. All cells were sorted at single-cell resolution, into a 384-well Eppendorf LoBind PCR plate (Eppendorf AG) containing 1 μL of lysis buffer (100 mM Triethylammonium bicarbonate (TEAB) pH 8.5, 20 % (v/v) 2,2,2-Trifluoroethanol (TFE)). Directly after sorting, plates were briefly spun, snap-frozen on dry ice for 5 min and then heated at 95 °C in a PCR machine (Applied Biosystems Veriti 384-well) for an additional 5 mins. Samples were then either subjected to further sample preparation or stored at -80 °C until further processing.

Bone marrow samples were collected from healthy donors following the standard protocol of the Department of Hematology, Rigshospitalet, with prior informed and written consent according to the Helsinki declaration under a protocol approved by the Danish National Ethics committee (1705391). Cell sorting was done on a FACS Symphony S6 instrument, controlled by the DIVA software package (v.8.0.2) and operated with a 100 μm nozzle. All cells were sorted at single-cell resolution, into a 384-well Eppendorf LoBind PCR plate (Eppendorf AG) containing 1 μL of lysis buffer (100 mM Triethylammonium bicarbonate (TEAB) pH 8.5, 20 % (v/v) 2,2,2-Trifluoroethanol (TFE)). Directly after sorting, plates were briefly spun, snap-frozen on dry ice for 5 min and then heated at 95 °C in a PCR machine (Applied Biosystems Veriti 384-well) for an additional 5 mins. Samples were then either subjected to further sample preparation or stored at -80 °C until further processing.

### Preparation of single cells for mass spectrometry analysis

Well plates containing single-cell protein lysates were digested with 2 ng of Trypsin (Sigma cat. nr. T6567) supplied in 1 μL of digestion buffer (100 mM TEAB pH 8.5). The digestion was carried out overnight at 37 °C, and subsequently stopped by the addition of 1 μL 1 % (v/v) trifluoroacetic acid (TFA). The resulting peptides were either directly submitted to mass spectrometry analysis or stored at -80 °C until further processing. All reagent dispensing was done using an I-DOT One instrument (Dispendix).

### Liquid chromatography

Chromatographic separation of peptides was conducted on a Vanquish Neo UHPLC system connected to a 50 cm □PAC Neo Low-load and an 10 □m EASY-spray emitter via built-in NanoViper fittings (all ThermoScientific). All separations were carried out with the column oven set to 50 °C. For comparison to Orbitrap Eclipse, FAIMS evaluation and 250pg method optimization and a 16 method with 3 minuted sample pickup overhead was used, where the percentage of buffer B (80 % ACN in H2O, 0.1 % FA) was initially increased from 4% to 8 % (0-0.2 min), 8% to 18% (0.2-2 min) with the nominal flow set at 750 nl/min, the flow was then reduced to 200nl/min (2 min-2.1 min) and the buffer B percentage increased to 18.1% (2.1 min 5.1min), 18.1% to 48% (5.1 min - 7.6 min), 48% to 99% (7.6min - 8.0min) and kept constant for 8min. For the MS1 and MS2 quantification comparisons and all single-cell runs trap-elution configuration. A 20min method with 2.5min autosampler overhead was utilized, where buffer B was increased from 4% to 10 % (0-0.2 min), 10% to 20% (0.2-3 min) with the nominal flow set at 750 nl/min, the flow was then reduced to 200nl/min (3 min-3.1 min) and the buffer B percentage increased to 28.1% (3.1 min 5.1min), 28.1% to 48% (5.1 min - 7.6 min), 48% to 99% (7.6min - 8.0min) and kept constant for 8min. Buffer B was then decreased to 1% for 2 min, after which the flow-rate was increased back to 750nl/min and kept for 2min. All Hela digest samples were injected from 96-well plate and all single-cell samples were directly injected from 384 well plate in which the samples were prepared as described.

### Mass spectrometry data acquisition

Acquisition of single-cell derived peptides was conducted with an Orbitrap Astral mass spectrometer operated in positive mode with the FAIMS Pro interface (Thermo Fisher Scientific) using a compensation voltage set to -48 V. Orbitrap MS1 spectra were acquired with the Orbitrap at a resolution of 240,000 and a scan range of 400 to 800 m/z with normalized automatic gain control (AGC) target of 300 % and maximum injection time of 100 or 200 ms depending on the method. Data independent acquisition of MS2 spectra was performed in the Astral using loop control set to 0.6 seconds per cycle with different static isolation window widths and injection times. For assessing FAIMS (Figure 2) the following widths and IITs were used: 3.4m/z - 3ms, 6.8m/z - 6ms, 13.6m/z - 12ms, 27.2 m/z - 24ms. For sensitivity optimization (Figure 3): 2.5 m/z - 5 ms, 5 m/z - 10 ms, 10 m/z - 20 ms, 14m/z - 30 ms, 20 m/z - 40 ms, 25 m/z - 50 ms, 30 m/z - 60 ms. For comparing MS1 and MS2 quantitative accuracy (Fiugre 4-6) IIT of 200 and 40 ms were used, for MS1 and MS2, isolation window width was set to 24 m/z. For samples with signal reminiscent of single-cell (Figure 5d-e) or true single-cells (Figure 7), 200 ms and 60 ms was used, with the isolation window set to 20 m/z. A 1 m/z window overlap and a range of 400-800 m/z was used for all methods. Fragmentation of precursor ions was performed using higher energy collisional dissociation (HCD) using a normalized collision energy (NCE) of 25 % and AGC target was set to 500 %.

### Mass spectrometry raw data analysis

All generated raw files were processed using Spectronaut version 18.4 and DIA-NN 1.8.1. For Spectronaut, direct DIA analysis was performed in pipeline mode using default BGS factory settings unless specified. The digestion enzyme was set to Trypsin/P and maximum 2 missed cleavages were allowed. The maximal number of modifications per peptide was set to 5. Protein N-terminal acetylation and methionine oxidation were set to variable and cysteine carbamidomethylation as fixed modification. For single-cell experiments carbamidomethylation was removed. DirectDIA+ (Deep) workflow was used with FDR on PSM, peptide and protein group set to 0.01. Quantification level was set to MS1 and the quantity type set to MaxLFQ. For DIA-NN Trypsin/P was set was the digestion enzyme with 1 missed cleavage allowed per peptide. N-terminal M excision and cysteine carbamidomethylation were included as modifications. The mass accuracy was determined automatically. Precursor level FDR was set to 1% and protein inferences was set to “Genes”, Heuristic protein inference on and MBR off. Use of isotopologues was on and “Robust LC (high-precision)” was used as the quantification strategy. Cross-run normalization was switched off.

### Data filtering and analysis

All exported report tables from both Spectronaut and DIA-NN were further analyzed with custom scripts based on python and visualized with R in the Visual Studio Code editor environment (see Code availability).

## Data availability

The raw proteomics data has been deposited to the following repository: MSV000095333.

## Code availability

The quantitative matrices used to generate all the figures in the manuscript are available in zenodo (10.5281/zenodo.12754964) and the analysis code available on Github (https://github.com/Schoof-Lab/Astral-scpMS)

## Author Contributions

V.P., P.A.F., and E.S. designed the study. V.P., P.A.F., and N.Ü. performed the experiments.

V.P., performed data analysis. V.P., P.A.F., B.F., T.N.A., E.Da., and E.M.S designed the methods and evaluated the data, with input from H.S., E.De., J.P., A. P., C.H, A.M., V.Z., and B.T.P. The manuscript was drafted and revised by V.P., P.A.F., J.W. and E.M.S., and has been read and approved by all authors. E.M.S. supervised the work.

## Acknowledgements

This work was funded by the following grants to E.M.S.: 1) reference number NNF21OC0071016 from the Novo Nordisk Foundation; 2) case no. 2067-00053B from the Independent Research Fund Denmark., 3) Leo Foundation (LF-OC-21-000832), and 4) Danish Cancer Society (R324-A17978) B.F. is the recipient of a fellowship from the Novo Nordisk Foundation as part of the Copenhagen Bioscience PhD. Programme, supported through grant NNF19SA0035442. Work in the Porse lab was supported by grants from the Svend Andersen Foundation, the Candys Foundation, the Danish Cancer Society, the Eva and Henry Frænkel Memorial Foundation, the Independent Research Fund Denmark and through a center grant from the Novo Nordisk Foundation (Novo Nordisk Foundation Center for Stem Cell Biology, DanStem; Grant Number NNF17CC0027852). This work has been performed in the context of the Danish Research Center for Precision Medicine in Blood Cancers funded by the Danish Cancer Society (R223-A13071) and Greater Copenhagen Health Science Partners.

